# Control of Septoria tritici blotch by winter wheat cultivar mixtures: Meta-analysis of 19 years of cultivar trials

**DOI:** 10.1101/658575

**Authors:** Rose Kristoffersen, Lise Nistrup Jørgensen, Lars Bonde Eriksen, Ghita Cordsen Nielsen, Lars Pødenphant Kiær

## Abstract

Wheat is the most commonly grown cereal crop in Europe and in major parts the most yield limiting disease is Septoria tritici blotch (STB). Currently, the control of the disease depends on cultivar resistance and significant input of fungicides. The impact of using mixtures of elite cultivars as an alternative was investigated through a meta-analysis based on trial data from the Danish national cultivar testing. The cultivar testing includes a four-way cultivar mixture every year and in these trials STB severity and yield have been monitored at multiple locations between 1995 and 2017. Results from 19 years of cultivar testing trials provided a data set for 406 trials from which the effect of mixtures was evaluated. The meta-analysis revealed that cultivar mixtures reduced STB severity with 10.6% and increased yields with 1.4% across all trials. The effects were greatest in untreated trials where STB severity was reduced with 17% and yields increased with 2.4%. The mixtures did not only perform better than the average of their component cultivars grown as pure stand, they also performed better than the average of the four most grown cultivars in a given year. No relationship was found between disease pressure or location and the performance of the mixtures. The mixtures included in the cultivar testing were not designed to control STB and the results are therefore perceived as a baseline to the attainable disease control from mixtures. The use of cultivar mixtures is relevant for low input farming systems, but can also contribute to disease control in intensive farming systems. Cultivar mixtures have the potential to minimise dependency on fungicides as an important element in integrated pest management.

## 1. Introduction

Septoria tritici blotch (STB) caused by the fungus *Zymoseptoria tritici* is among the most devastating wheat diseases in Europe (Fones and Gurr, 2015). In the European Union (EU28), agriculture utilises 175 hectares, equivalent to 40% of the total area (Eurostat, 2015). According to The FAO (2013) wheat alone accounts for 33% of this area (Food and Agriculture Organization of the United Nations, 2019). The intensive wheat cultivation and high rainfall in the major wheat producing countries in Western Europe provides favourable conditions for a splash-dispersed disease as STB (Torriani et al., 2015). Without control measures, yield losses can reach 10-30% annually (Jørgensen et al., 2014). Currently, STB control is dominated by azole and SDHI fungicides (Kirikyali et al., 2017; Torriani et al., 2015). A pronounced reduction in field performances using azoles has been seen in many parts of Europe (Jørgensen et al., 2017) and also SDHIs are now challenged due to starting resistance in countries like Ireland and UK (Kildea et al., 2016). Alternative disease management strategies are necessary.

The annual disease epidemics in cereals are specific to cultivated ecosystems and are not found in their natural counterparts (McCann, 2000). Modern cropping systems with genetically uniform crops are very vulnerable to biotic and abiotic disturbances and make an optimal environment for the formation and spread of pathogens (McCann, 2000; McDonald and Stukenbrock, 2016). Increasing biodiversity in cropping systems has been proposed as a key factor in creating more robust systems with less need for chemical control and essential in an integrated pest management framework (Stenberg, 2017). Diverse systems as intercropping or cultivar mixtures is associated with subsistence agriculture, but has been investigated in more modern agriculture (Smithson and Lenné, 1996). Lower forms of diversity such as cultivar mixtures that are diverse on gene level has great potential in reducing the negative effects of monocultures and increasing the production (Barot et al., 2017). Cultivar mixtures can reduce the epidemic development of a disease and increase yields (Borg et al., 2018; Finckh et al., 2000; Wolfe, 1985). Diversification has clear benefits, but when implementing methods that are not part of current farming practice it is important to address concerns from the growers about yields and management. Cultivar mixtures are low hanging fruit in increasing genetic diversity in the cereal production as the implementation requires little disruption of the management (Finckh et al., 2000).

Studies with mixtures specifically designed to control STB have showed potential in reducing this disease (Cowger and Mundt, 2002; Gigot et al., 2013; Mille et al., 2006; Vidal et al., 2017a). Meta-analyses of published articles have found diseases to play an important role in achieving yield increases through mixtures. Kiær et al. (2009) found a significant increase in yield by mixing cultivars when the component cultivars differed in disease resistance. Borg et al. (2018) documented a general yield increase by mixtures of 3.5% and 6.2% when mixtures were grown under high disease pressure. The meta-analysis by Reiss and Drinkwater (2018) found mixtures that were designed based on both disease and physical traits to increase yield under high disease pressure.

To investigate the effect of cultivar mixtures on STB a meta-analysis of field trial data from the national cultivar testing was conducted. Since 1991, official Danish cultivar trials have included a cultivar mixture as the standard against which other cultivars are measured. The cultivar mixture was introduced to ensure a stable measurement basis and to increase the continuity in the trials (Pedersen, 1995). These trials include random cultivar mixtures of high yielding cultivars available to the farmers and were not designed for STB control. As the purpose of the mixture design can have an impact on the efficacy of the mixture (Borg et al., 2018; Reiss and Drinkwater, 2018) the results from the non-designed mixtures in this study are produced with the aim of estimating a baseline for STB control by mixtures of elite cultivars. Even though the trials were not designed they still include mixtures with different proportions of susceptible and resistant cultivars and this information can be used to predict how mixtures can be designed to reduce STB. To make the obtained results applicable to farming in practice the efficacy of the mixtures were evaluated both on disease severity and on yield. A disease reduction is irrelevant to the farmer if mixtures in general are accompanied by a yield reduction. The performance of the mixtures was not only compared to the component cultivars but also to the cultivars the farmers commonly grow to evaluate the performance of the mixture against the standard practice.

The hypotheses tested in this study: 1) Non-designed cultivar mixtures of elite cultivars reduce STB severity and increase yields compared to the average of component cultivars, 2) The mixtures reduce STB severity and increase yield compared to the most grown cultivar in a given season, 3) Lower STB severity in mixtures contributes to higher yields in mixtures, 4) STB score of components from the previous season can predict the effect of cultivar mixing on STB disease severity.

## 2. Materials and methods

### 2.1. Cultivar trials 1995-2017

Cultivar testing has been performed in Denmark from 1995-2017 in a collaboration between SEGES (the Farmers Union), and Tystoftefonden, which is performing Value for Cultivation and Use (VCU) tests for the national listing. During this period, field trials were conducted using a similar protocol across sites and years. A mixture of four cultivars was defined each year to be used as a reference. Original observation data from these trials was used in this study to investigate the effects of these non-designed cultivar mixtures of winter wheat on STB severity and yield.

The winter wheat cultivars included in the reference cultivar mixture have been continuously up for evaluation replacing a single component cultivar each year (table 1) (Pedersen, 1995). 73% of the mixtures included cultivars from three different breeders (not shown). Data regarding the most commonly grown cultivars were obtained from sales statistics at sortinfo.dk and included for comparison (table 1).

**Table 1:** The components of the standard cultivar mixture included in the national Danish cultivar tests 1995-2017 and the cultivars with the largest certified seed sale each year (SortInfo.dk). *Italics* denote new cultivars in the reference mixture. Use statistics (and thus the most grown cultivars) was first registered in 1999.

| Year | Mixture components | Most grown cultivars in given year |
| --- | --- | --- |
| 1994 | Pepital, Haven, Hereward, Nova | - |
| 1995 | Pepital, Haven, Hereward, *Hussar* | - |
| 1996 | *Ritmo*, Haven, Hereward, Hussar | - |
| 1997 | Ritmo, *Trintella*, Hereward, Hussar | - |
| 1998 | Ritmo, Trintella, *Pentium*, Hussar | - |
| 1999 | Ritmo, Trintella, Pentium, *Cortez* | Ritmo, Stakado, Trintella, Lynx |
| 2000 | Ritmo, Trintella, Pentium, Cortez | Ritmo, Stakado, Kris, Cortez |
| 2001 | Ritmo, *Terra*, Pentium, Cortez | Ritmo, Kris, Stakado, Baltimor |
| 2002* | Ritmo, *Solist*, Pentium, Cortez | Bill, Ritmo, Stakado, Kris |
| 2003 | Ritmo, Solist, Pentium, *Boston* | Solist, Bil, Senat, Ritmo |
| 2004* | Ritmo, Solist, *Galicja*, Boston | Deben, Grommit, Hattrick, Bill |
| 2005* | Ritmo, Solist, Galicja, *Skalmeje* | Deben, Hattrick, Robigus, Smuggler |
| 2006 | Ritmo, Solist, *Ambition*, Skalmeje | Smuggler, Robigus, Hattrick, Samyl |
| 2007 | *Fru ment*, Solist, Ambition, Skalmeje | Smuggler, Skalmeje, Robigus, Samyl |
| 2008 | Fru ment, *Hereford*, Ambition, Skalmeje | Ambition, Smuggler, Skalmeje, Fru ment |
| 2009 | Fru ment, Hereford, Ambition, *Contact* | Fru ment, Ambition, Hereford, Smuggler |
| 2010 | Fru ment, Hereford, Ambition, *Mariboss* | Hereford, Fru ment, Ambition, Oakly |
| 2011 | Fru ment, Hereford, *Jensen*, Mariboss | Hereford, Mariboss, Fru ment, Tabasco |
| 2012 | *KWS Dacanto*, Hereford, Jensen, Mariboss | Hereford, Mariboss, Jensen, Tabasco |
| 2013 | KWS Dacanto, Hereford, Jensen, Mariboss | Mariboss, Jensen, Hereford, KWS Dacanto |
| 2014 | KWS Dacanto, Hereford, Jensen, Mariboss | Mariboss, Jensen, KWS Dacanto, Hereford |
| 2015 | KWS Dacanto, *Benchmark*, Jensen, Mariboss | Mariboss, KWS Dacanto, Hereford, Jensen |
| 2016 | KWS Dacanto, Benchmark, *Torp*, Mariboss | Torp, Mariboss, KWS Dacanto, Hereford |
| 2017 | KWS Dacanto, Benchmark, Torp, *Kalmar* | Torp, Benchmark, Pistoria, KWS Lili |

\* One or more component cultivars missing in pure stand, not included in the analysis

The trials were sown as randomised complete block design or alpha-design with incomplete blocks with three to six replicates. Plot sizes varied from 14 m^2^ to 20 m^2^ placed at locations across Denmark at farmers’ fields, field stations and breeding companies. Thy were either one-factor trials with only fungicide-treated plots, or two-factor trials including both treated and untreated control plots. Fungicide treatments were applied by the individual farmers and varied between locations according to local practices, consisting of two to three treatments with a reduced dose of the most effective azoles, strobilurins and SDHIs at the time. Typically, the dose rate applied was 30-75% of a full standard rate for the compound. Additionally, fungicide treated and control plots in each trial were all treated with herbicides, insecticides, growth regulators and fertiliser according to the requirement at the individual location. The harvested grain was measured as kg/plot and and transferred to yield per ha (dt/ha) adjusted to 15% moisture content. STB severity was assessed visually around GS 69-75 two weeks after the last fungicide treatment of the trial, typically around applied in the period 5.-20. June. Severity was measured as a total plot score of percentage disease coverage by technicians from the different trial units.

### 2.2. Data filtering and meta-analysis

#### 2.2.1. Filtering of data

Data quality was ensured by filtering data according to a number of criteria. Only field trials where all four component cultivars were also grown in pure stand were included, resulting in the exclusion of three trial years (2002, 2004 and 2005). Only trials with data from at least three replicates, and STB severity data from at least half the plots, where included in the analysis. For trials with more than two fungicide treatments (e.g. spraying rates; 30 trials in total), only control (untreated) and standard treatments were used. To avoid unrealistically large or small values when calculating STB mixing effects (see below), a minimum threshold of 2% STB disease severity (averaged across the four components) was applied. From the data obtained, yield was registered for all trials, but STB seveity data was sometimes absent.

#### 2.2.2. Process

The data analysis consisted of three steps:

1. Calculation of yield and STB severity estimates for the cultivars and reference mixture in each trial.
2. Calculation of mixing effect for each trial.
3. Meta estimates for yield and STB mixing effects through meta-analysis.

#### 2.2.3. Estimating yield and STB severity in field trials

Yield and STB estimates for the cultivars and reference mixture were obtained for each field trial using linear mixed models per the package ‘lme4’ (Bates et al., 2015) for R (R Core Team, 2018). Models were fitted for the different designs included in the experiments Models were fitted for the different designs included in the experiments as obs ~ ct + *block* | *rep* + *rep*, where obs denotes grain yield measurements or STB disease severity (as percentages), ct denotes the cultivar (pure stands and mixture) used as explanatory factor, *block* denotes the random intercepts of incomplete blocks (for alpha designs only), and *rep* denotes the random intercepts of replicates. A subset of the data was tested and found to follow a normal distribution for yield and a log-normal distribution for STB and so the remaining STB data was log transformed before further analysis.

#### 2.2.4. Mixing effects

Estimates from the fitted linear mixed models with yield and STB as response variables, respectively, were used to calculate mixing effects of either response type, using the log response ratio (lnRR) as described by Hedges et al. (1999). The lnRR is the natural logarithm of the ratio between treatment (here, the mixture estimate) over the control (here, the average of the pure stand estimates of component cultivars).

#### 2.2.5. Meta-analysis and moderators

Meta-analysis was performed by fitting mixed models as integrated in the rma.mv function of the R package ‘metafor’ (Viechtbauer, 2010). Effect sizes lnRR and their variances were used as response variable, and year and year-within-location were used as random effects. A series of meta-analyses were performed with different moderator variables (table 2) that might influence mixing effect.

**Table 2:** List of moderators used and the various mixing effects they have effect on.

| Effect of | On |
| --- | --- |
| STB mixing effect | Yield mixing effect |
| Disease pressure | STB and yield mixing effect |
| Component susceptibility | STB mixing effect |

Disease pressure was estimated for each field trial in two ways. First, as the average STB severity of all pure stand cultivars in the trial in treated or untreated plots. Second, as average yield response to fungicide treatment, calculated per location and based on the fungicide response map created by SEGES and the Danish Technological Institute (DTI) (figure 1) as calculated from field trial data in the period 2002-2016 (Nielsen and Trénel, 2017). This local fungicide response was used as an indirect measure of disease pressure as estimates were based on trial data on response from diseases. Septoria was the main target for the modelling and yellow rust (*Puccinia striiformis*) above 1% was excluded from the analysis (Nielsen and Trénel, 2017). These estimates of disease pressure (1-10) were linked to each trial location by postcode (local disease pressure). The map was further divided into seven regions with similar scores (regional disease pressure). ‘High disease pressure’ was defined as disease pressure above 8%, with 66% of data for treated plots falling below this threshold.

**Figure 1:**
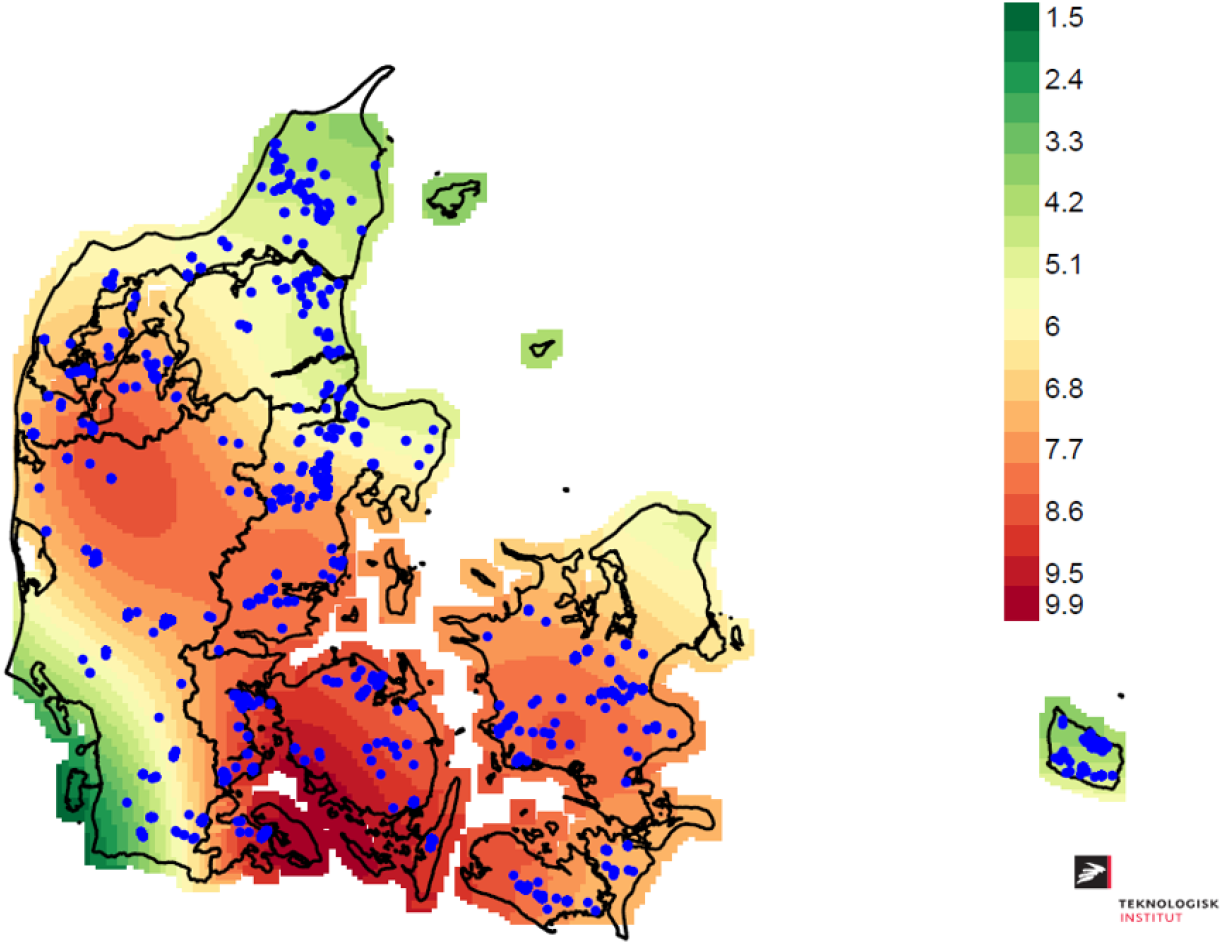
Map of yield increase (hkg/ha) from fungicide applications used to estimate disease pressure. Blue dots represent trial locations. The map was produced by SEGES and the Danish Technological Institute (DTI) (Nielsen and Trénel, 2017).

Component susceptibility information was obtained from the official susceptibility score of the individual component cultivars that was measured in the national observation plots (published on SortInfo.dk) and available at the time of sowing each year. These scores were used to derive single moderators including means, ranges and standard deviations, or categorised into susceptible/resistant based on different thresholds for a resistant cultivar.

Meta-estimates were back transformed (as exp(estimate)-1) in order to report the relative difference between mixture and pure stand yields and disease observations.

## 3. Results

The filtering of field trial data resulted in 569 unique mixtures cultivated across 406 trials. Of these, 319 mixtures across 221 trials could be used to estimate the overall STB mixing effect. Treated and untreated plots differed in STB severity and yields (table 3). On average untreated plot were more affected by STB and had lower yields compared to the fungicide treated plots. There was a significant negative correlation (p < 0.001) between STB severity and yield across the trials with an estimated effect of 19% meaning that field with a disease severity of 50% would correspond to a 9.5% yield loss. The susceptibility to STB for specific cultivars included in the mixtures ranged from 0.8% to 19% with an average of 7.7% when grown in pure stand.

**Table 3:** Average yield and disease severity across the field trials included. In parenthesis the standard deviation of the estimate.

|  | STB disease severity, % | Yield, hkg/ha |
| --- | --- | --- |
| Untreated plots | 16.3 ($\pm 16.8$) | 78.5 ($\pm 15.9$) |
| Treated plots | 6.1 ($\pm 7.9$) | 88.8 ($\pm 15.2$) |

### 3.1. Mixing effects on STB

Across all field trials, the non-designed cultivar mixtures reduced STB severity with 10.6% compared to the average of the component cultivars in pure stand (figure 2). The highest reduction was observed in untreated plots where mixtures reduced STB severity with 17%. Mixtures exhibited 12.3% less STB than the average of the four most grown cultivars each year, overall (figure 2). This effect was greater for treated than for untreated plots, meaning that in treated plots the STB reduction from mixing was larger than in untreated plots. When comparing mixtures with the single most grown cultivar, reductions were similar although there were no differences between treated and untreated plots (data not shown). Disease reduction was significant in all cultivar mixtures shown in fig 2 (all p < 0.001).

**Figure 2:**
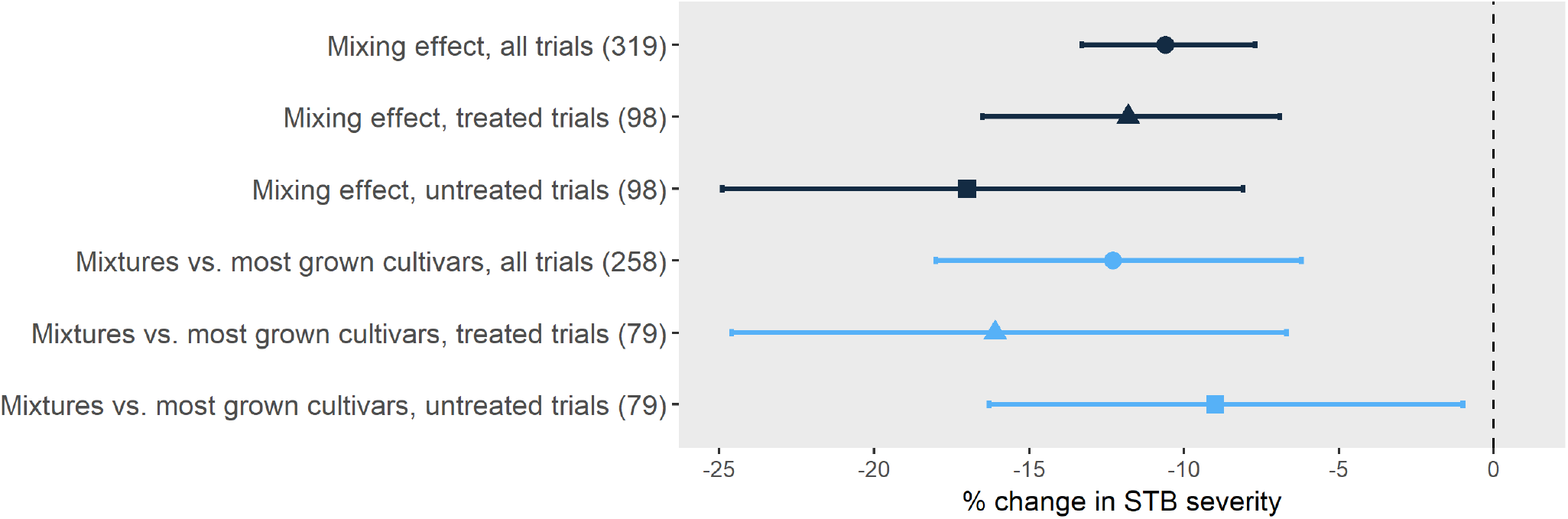
Mixing effects measured as % change in STB severity. The upper three effects (black lines) are mixing effects relative to the average of the component cultivars. The lower three effects (blue lines) are STB of mixtures relative to the average of the four most grown cultivars the given year. The effects were evaluated both across all trials (•) and separately for treated (▲) and untreated trials (■). Mean values and 95% confidence intervals are shown. In parenthesis is the number of mixture data points for each effect, *k*.

#### 3.1.1. Disease pressure

The reduction in STB severity in mixtures was expected to increase with increasing disease pressure. Against expectations, this could not be confirmed regardless of the disease pressure was measured in treated or untreated plots. The effect sizes were numerically small and non-significant (data not shown). Measuring disease pressure indirectly via the estimated yield increase from fungicide applications at the trial site also did not show an effect on the STB mixing effect. Expanding this to broader disease pressure regions yielded numerically higher effect sizes, but with equally high variation and lack of significance (data not shown).

#### 3.1.2. Mixture composition

The effect of mixture composition with regards to STB susceptibility was evaluated based on the mean (component susceptibility average) or the variability (measured as range or standard deviation) in component susceptibility. However, neither of these were found to affect the STB mixing effect.

Dividing components into categories of either ‘susceptible’ or ‘resistant’ showed an impact on STB mixing effect (figure 3). If the threshold for a resistant cultivar was set to either 4% or 5% STB mixtures with one (25%) resistant cultivar were most successful in reducing STB severity, as compared to the average of component cultivars in pure stand. If the threshold was set to 6% or 7% STB, there was no difference between mixtures compositions. The 4% threshold was the only case where a mixture exceeded 20% STB reduction. Only threshold 4% and 7% were included in figure 3. Given that the mixtures were not designed to include specific proportions of susceptible and resistant cultivars, the number of mixtures in each category varied between thresholds (table 4). If the threshold was set to 5% STB there was eight mixtures with no resistant cultivars and only two mixtures with two resistant cultivars. The composition with the highest reduction (4% STB susceptibility, one resistant cultivar) is only represented by one mixture.

**Figure 3:**
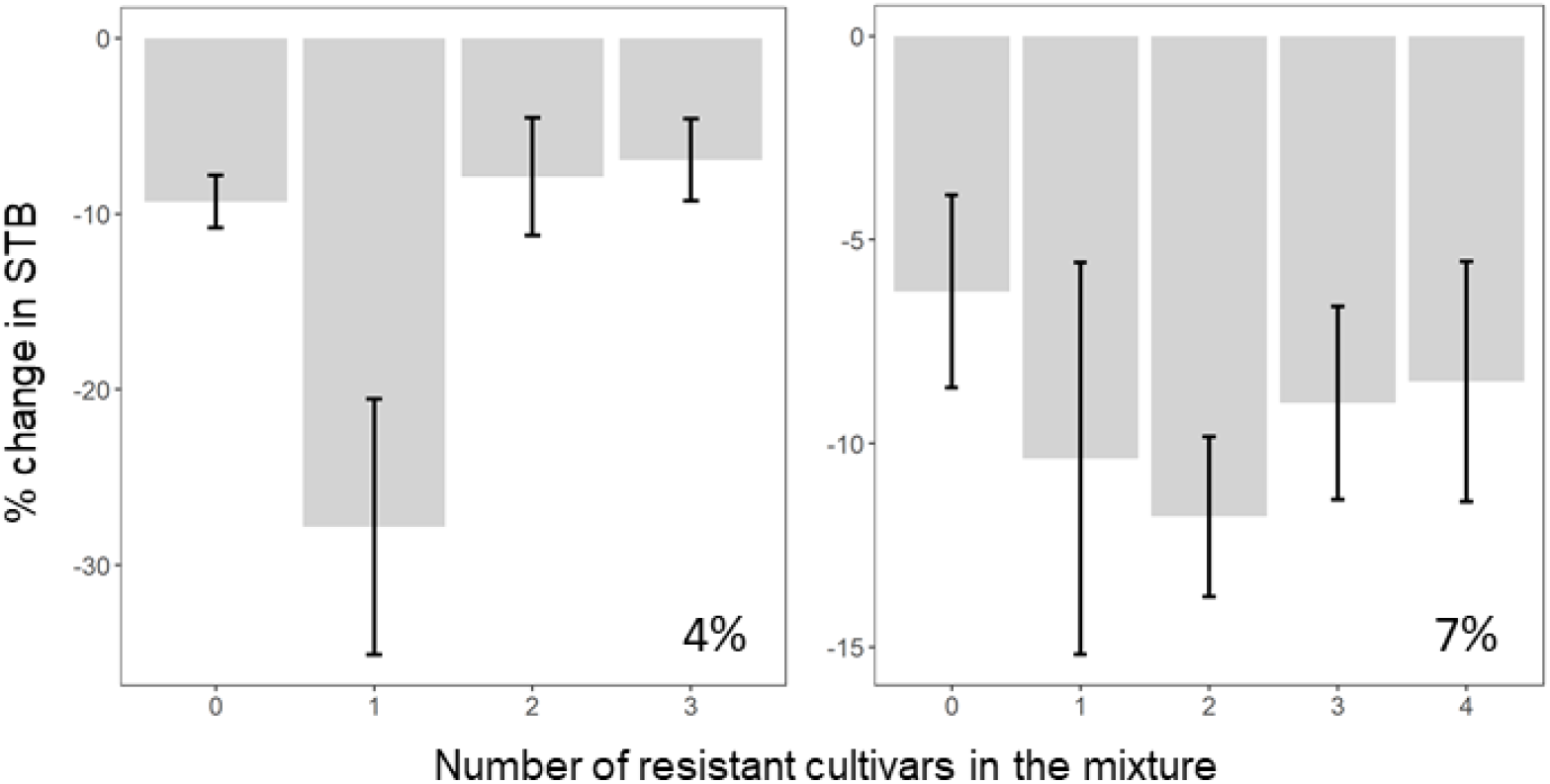
Effect of the ratio of resistant/susceptible component cultivars on STB mixing effect. Threshold of what constitutes a resistant cultivar shown for <4% STB (left) and <7% STB (right). X-axis denotes number of resistant cultivars in the mixture, y-axis denotes STB mixing effect. Bars represent mean effect and error bars +/− standard deviation.

**Table 4:** Number of cultivar mixtures available in the data set for mixtures with different proportions of resistant cultivars. The number of mixtures varied depending on the threshold for what defines a resistant cultivar. The resistance threshold was set to different values of STB severity measured as % disease cover in treated plots.

| Resistance threshold (%STB) | Resistant cultivars in the mixture |  |  |  |  |
| --- | --- | --- | --- | --- | --- |
|  | None | One | Two | Three | Four |
| 4% | 11 | 1 | 3 | 3 | 0 |
| 5% | 8 | 4 | 2 | 3 | 1 |
| 6% | 7 | 4 | 2 | 4 | 1 |
| 7% | 5 | 1 | 6 | 4 | 2 |

### 3.2. Mixing effect on grain yield

Mixtures gave an overall grain yield increase of 1.4% compared to the average of the component cultivars (figure 4). The effect was greater in untreated plots showing a yield increase of 2.4%. Mixtures also increased yield with 1.4% compared to an average of the four most grown cultivars a given year (figure 4). The yield increase was equally greater for untreated plots with a yield increase of 2.9%. The findings were similar when comparing mixtures with only the single most grown cultivar (data not shown). All yield increases were significant (p < 0.001) except for the comparison with the most grown cultivars in treated plots (p 0.5). The greater yield increases in untreated plots could indicate a correlation between STB reductions and yield increases in mixtures. however, such a correlation could not be confirmed (figure 5). This was true when looking at trials overall, at untreated trials only and when looking specifically under high disease pressure.

**Figure 4:**
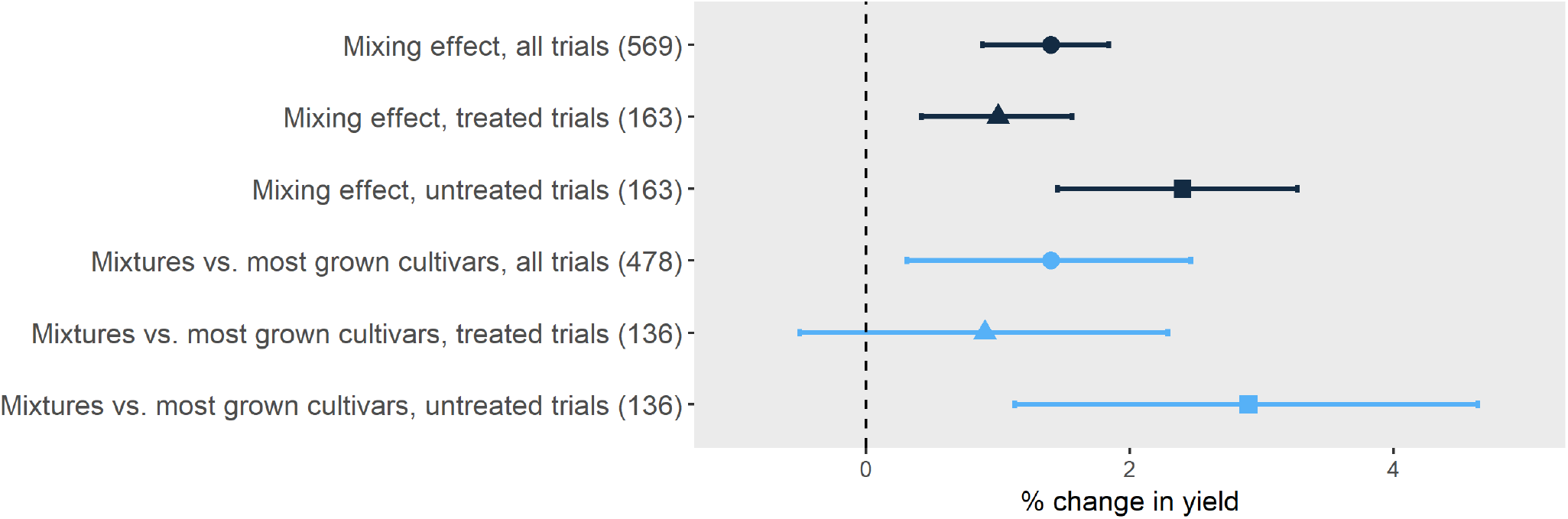
Mixing effects measured as % change in yield. The upper three effects (black lines) are mixing effects relative to the average of the component cultivars. The lower three effects (blue lines) are yield of mixtures relative to the average of the four most grown cultivars the given year. The effects were evaluated both across all field trials (•) and separately for treated (▲) and untreated trials (■). In parenthesis is the number of mixture data points for each effect, *k*. Mean values and 95% confidence intervals are shown.

**Figure 5:**
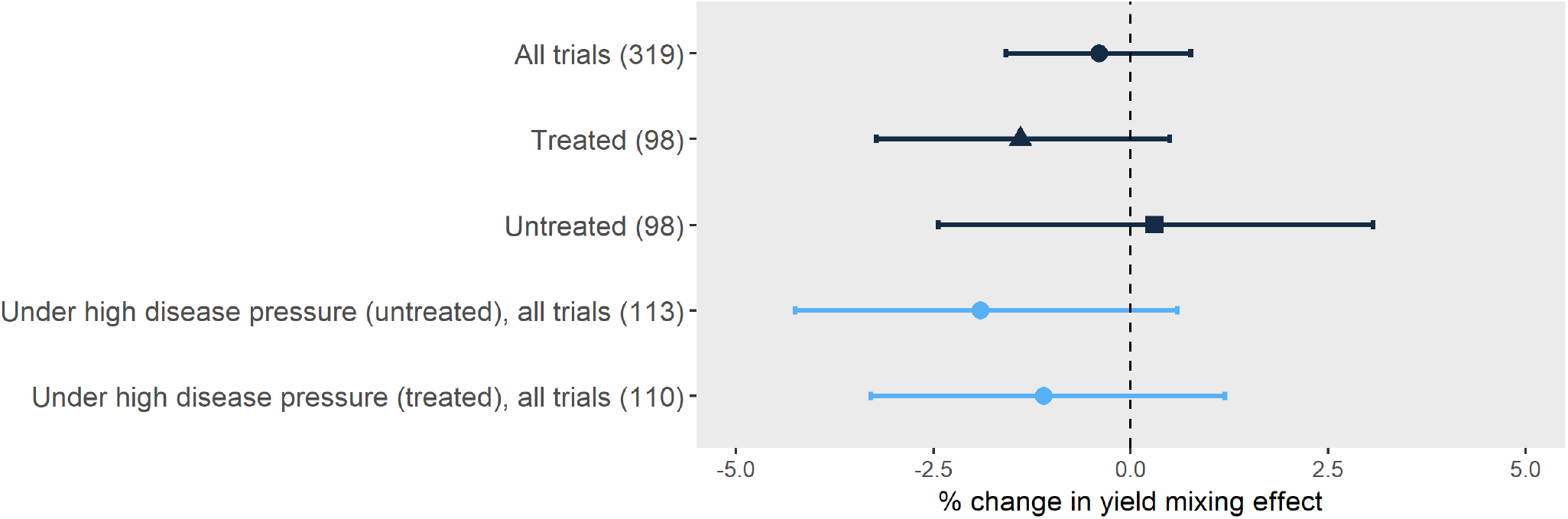
Effect of STB mixing effect on yield mixing effect (% change). The upper three effects (black lines) were evaluated both across all field trials (•) and separately for treated (▲) and untreated trials (■). The lower two effects (blue lines) are based on trials with high disease pressure (mean STB of all pure stand cultivars >8%), measured from untreated and treated plots, respectively. In parenthesis is the number of mixture data points for each effect, *k*. Mean values and 95% confidence intervals are shown.

## 4. Conclusion and discussion

### 4.1. Mixtures reduce STB and increase yields

The cultivar mixtures in this study were successful at reducing STB and increasing yields compared with their pure stand counterparts. This was in line with the overall positive results obtained in the previous mixtures studies conducted with STB (Cowger and Mundt, 2002; Gigot et al., 2013; Mille et al., 2006; Vidal et al., 2017a) and the extensive studies documenting yield increases from mixtures (Kiær et al., 2009; Borg et al., 2018; Reiss and Drinkwater, 2018). It has, however, not previously been documented that cultivar mixtures performed better than the average of the most grown cultivars in a given season. The study found significant mixing effects and supported hypotheses 1) and 2): Non-designed cultivar mixtures of elite cultivars reduced STB severity and increased yields compared to the average of component cultivars and the mixtures reduced STB severity and increased yield compared to the most grown cultivar in a given season.

### 4.2. The impact of disease pressure

STB mixing effect has been reported to vary with disease pressure (Cowger et al., 2000; Gigot et al., 2013) and the lack of correlation between these factors shown in this study was unexpected. The absence of impact from disease pressure could be explained for a large part by the field trial data being in the lower range of disease pressure, leaving few trials with high disease pressure to observe any significant differences. It appears to be to common practice in the studied field trials to avoid mixing naturally senescent leaf area with diseased area at late growth stages by only assessing disease on green leaves (Ghita Cordsen Nielsen, personal communication). This practice will tend to underestimate the disease pressure especially in the most susceptible cultivars. Even though a guideline exists, the disease assessments were carried out by different persons which could have influenced the disease severity scale. This is not an issue for the mixing effect itself as it is relative to components from the same trial, assessed by the same person throughout the season. The second analysis with indirect disease pressure calculated from the fungicide-yield response map had the potential to provide a more stable and objective measurement of disease pressure. However, this indirect disease pressure did not correlate with STB mixing effect either. The weakness of using the fungicide response is that it cannot be quantified how much of the response was caused by the STB severity. Even though STB is considered the most yield limiting disease other diseases might have influenced the results at certain locations. The study could not provide an explanation to the variation in the mixing effects based on disease pressure.

### 4.3. Correlations between STB and yield

Diseases are among the factors to have the highest impact on yield mixing effect (Borg et al., 2018). It was not evident that the reduction in STB from mixtures was the sole cause of the yield increase in this study and a significant yield increase would not be expected from just a 10% STB reduction. However, as both the STB and yield effects were larger in untreated trials it could indicate a correlation. STB is the most important disease in Danish winter wheat production (Jørgensen et al., 2014) and was the only disease included in this study. Other diseases that are of less importance could theoretically affect the results and mixing effects are well-documented for wind borne diseases such as rust diseases (*Pucciniales*) and powdery mildew (*Blumeria graminis*) (Finckh et al., 2000; Huang et al., 2012; Newton et al., 2002). Other factors can contribute to the yield mixing effect as well such as nutrient utilization, canopy cover, winter hardiness, proneness to lodging etc. (Bessler et al., 2009; Bowden et al., 1980; Jackson and Wennig, 1997; Sarandon and Sarandon, 2006; Vidal et al., 2017b). It cannot be excluded that these other factors have had an impact on the yield mixing effect found in this study. However, as the only difference between treated and untreated trials was the fungicide application, it would be unlikely that abiotic factors would contribute differently to the yield mixing effects. Hypothesis 3) could not be confirmed. The results did not directly support that lower STB severity in mixtures contributed to higher yields in the mixtures. Further investigations are needed to explain the effect difference between treated and untreated trials.

### 4.4. Predicting mixing effects from components

The results provided no clear answers as to how mixtures should be designed. The components in the mixtures were not selected based on susceptibility to STB and can thus only indirectly be used to evaluate mixture design. In this study, neither high variation in component susceptibility or mixtures with overall very susceptible or very resistant components correlated with mixing effects on STB or yiels. A common approach in mixture studies is to compare mixtures with different ratios of susceptible and resistant cultivars (Wolfe, 1985) and analysing composition in this way had an impact on the mixing effects. However, in the current study the impact varied greatly depending on where the threshold for a resistant cultivar was placed. Based on this analysis a cultivar mixture with only one cultivar with less than 4% STB performed best. As there was only one mixture in the study that matched this criterion the result would need further validation. Increasing the threshold to 5% gave four mixtures with one resistant component, but did not provide as high a reduction as with the 4% threshold. The mixtures in this study were not designed to evaluate the effect of the ratio of resistant/susceptible cultivars and primarily contain cultivars that are moderately resistant to STB. The susceptibility difference in designed mixtures could be much higher than in this study. Some caution should be taken before applying the results from this study directly to conclude on mixture design as the components of the mixtures in the cultivar testing are changed every year and the performance to a specific mixture composition and the effect of the year cannot be excluded from the mixture performance. We found only partial support for hypothesis 4) that STB score of components from the previous season would be able to predict STB mixing effects.

### 4.5. Relevance of cultivar mixtures in practice

There is a need to change disease management to include a strategy where the cultivar genetics are utilized more efficiently for disease control. The use of cultivar mixtures is an easy tool that is ready to be implemented. For growers there is an incentive to grow mixtures as they both perform better than the average of the pure stand cultivars and better than the cultivars the farmers would otherwise choose. An argument often used against mixtures is that they do not perform better than the most resistant or highest yielding cultivar. This argument does not take into account that it is hard to predict which cultivar performs best the coming season. To reduce the risk of choosing the wrong cultivar a farmer might sow more than one cultivar on their farm and could reduce the risk further by sowing these cultivars in mixture. The reductions in STB severity observed in this study were not large enough to provide a significant disease control alone. If the results with non-designed mixtures can be viewed as a baseline of STB control by cultivar mixtures, mixtures specifically designed to control STB can be expected to yield higher and this could improve disease control and play a more important part in integrated pest management. Breeders may argue that instead of using cultivar mixtures, higher levels of resistance or more durable resistance to diseases should be achieved through pyramiding of genes in individual cultivars. This method has, however, not unequivocally solved issues with disease resistance in breeding (Grimmer et al., 2015) and cannot replace the mechanisms of cultivar mixtures. The ability of cultivar mixtures to suppress disease is commonly ascribed to the spatial distribution of resistance genes (Borg et al., 2018; Gigot et al., 2014; Newton and Guy, 2011). Application of mixtures in real farm settings with larger field sizes (Finckh et al., 2000) and more patchy distributions of the cultivars (Newton and Guy, 2009) are likely to contribute to larger effects as well. A criterion for cultivar mixtures to be relevant in practical agriculture was that they did not have any negative impacts on yield and they did result in a statistically significant yield increase. The yield increase of 1.5-2.5% might not be large but it is, as mentioned by Reiss and Drinkwater (2018), on level with the yield increase achieved through breeding. Some of the main barriers for wider use of cultivar mixtures are farming traditions and lack of incentive from seed companies when mixing components from different breeders including issues with royalties. The use of cultivar mixtures is especially well suited for farm types that predominantly use cereals for animal feed as is the typical scenario in Denmark (Lundø, 2017). It is more challenging to implement cultivar mixtures in malting barley and bread wheat production where the maltsters and millers typically require cultivars grown as monocultures (Finckh et al., 2000).

The results provide a solid basis for implementing cultivar mixtures in the wheat production. However, in order to optimise mixture design and improve performance more thoroughly designed trials are needed.

## 5. Acknowledgements

The authors wish to thank the Danish Technological Institute for providing the trial data and Thies Marten Heick (Aarhus University) and Jeanine Raw (University of Salford) for proofreading the manuscript.

